# HEx: a heterologous expression platform for the discovery of fungal natural products

**DOI:** 10.1101/247940

**Authors:** Colin JB Harvey, Mancheng Tang, Ulrich Schlecht, Joe Horecka, Curt R Fischer, Hsiao-ching Lin, Jian Li, Brian Naughton, James Cherry, Molly Miranda, Yong Fuga Li, Angela M Chu, James R Hennessy, Gergana A Vandova, Diane Inglis, Raeka Aiyar, Lars M. Steinmetz, Ronald W. Davis, Marnix H. Medema, Elizabeth Sattely, Chaitan Khosla, Robert P St.Onge, Yi Tang, Maureen E. Hillenmeyer

## Abstract

For decades, fungi have been a source of FDA-approved natural products such as penicillin, cyclosporine, and the statins. Recent breakthroughs in DNA sequencing suggest that millions of fungal species exist on Earth with each genome encoding pathways capable of generating as many as dozens of natural products. However, the majority of encoded molecules are difficult or impossible to access because the organisms are uncultivable or the genes are transcriptionally silent. To overcome this bottleneck in natural product discovery, we developed the HEx (Heterologous EXpression) synthetic biology platform for rapid, scalable expression of fungal biosynthetic genes and their encoded metabolites in *Saccharomyces cerevisiae*. We applied this platform to 41 fungal biosynthetic gene clusters from diverse fungal species from around the world, 22 of which produced detectable compounds. These included novel compounds with unexpected biosynthetic origins, particularly from poorly studied species. This result establishes the HEx platform for rapid discovery of natural products from any fungal species, even those that are uncultivable, and opens the door to discovery of the next generation of natural products.

**Summary:** Here we present the largest scale effort reported to date toward the complete refactoring and heterologous expression of fungal biosynthetic gene clusters utilizing HEx, a novel synthetic biology platform.

## Main Text

Natural products are indispensable to modern medicine, with 73% of antibiotics, 49% of anticancer compounds, and 32% of all new drugs approved by the FDA between 1980 and 2012 being natural products or derivatives thereof (*1*). Fungi are prolific producers of therapeutically relevant natural products(*2*, *3*), having yielded penicillin, the first widely used antibiotic; cyclosporine, the immunosuppressant that enabled widespread organ transplantation; and lovastatin, the progenitor of the statin class of cholesterol-lowering drugs. In all of these examples, compounds were isolated from laboratory cultures of single fungal isolates. Recent advances in genome sequencing have revealed that more than 5 million fungal species likely exist on the earth(*4*) with each species encoding as many as 80 natural product biosynthetic pathways (*5*, *6*). However, despite the increased ease of DNA sequencing, fungal cultivation remains a bottleneck: only a fraction of the fungi in any given environmental sample have been cultured under laboratory conditions(*7*). Even within cultured species, the majority of biosynthetic gene clusters (BGCs) present in the genome are either transcriptionally silent or expressed at very low levels(*8*). The identification and expression of these BGCs thus presents a major opportunity for the discovery of novel natural products.

Previous approaches for surveying transcriptionally silent, or cryptic, fungal BGCs for the production of novel compounds have included BGC activation within the native host through promoter or transcription factor manipulation(*9*–*11*), CRISPR-based genome editing(*12*), and epigenetic activation(*13*–*15*). These approaches, however, are limited to those BGCs whose native hosts are both culturable and genetically tractable. For cryptic BGCs within the genomes of several *Aspergilli*, heterologous expression by cloning of large intact contigs into *Aspergillus nidulans* has yielded several new natural products(*16*).

Heterologous expression by complete BGC refactoring is an approach that is agnostic to the native host of a BGC, permitting access to cryptic BGCs from potentially any organism(*17*–*19*). We present HEx, an improved, scalable approach to heterologous expression of cryptic fungal BGCs(Figure 1). HEx comprises bioinformatic tools to identify and prioritize BGCs in genome data, genetic parts to refactor BGCs for expression in *S. cerevisiae*, background strains with improved growth and expression phenotypes, and synthetic biology tools to assemble and express synthetic DNA in the heterologous host (Figure 2-4). Strains expressing BGCs were analyzed via untargeted metabolomics(*20*), and if the compound appeared novel, full structures were solved using LC-MS and NMR.

**Figure 1:**
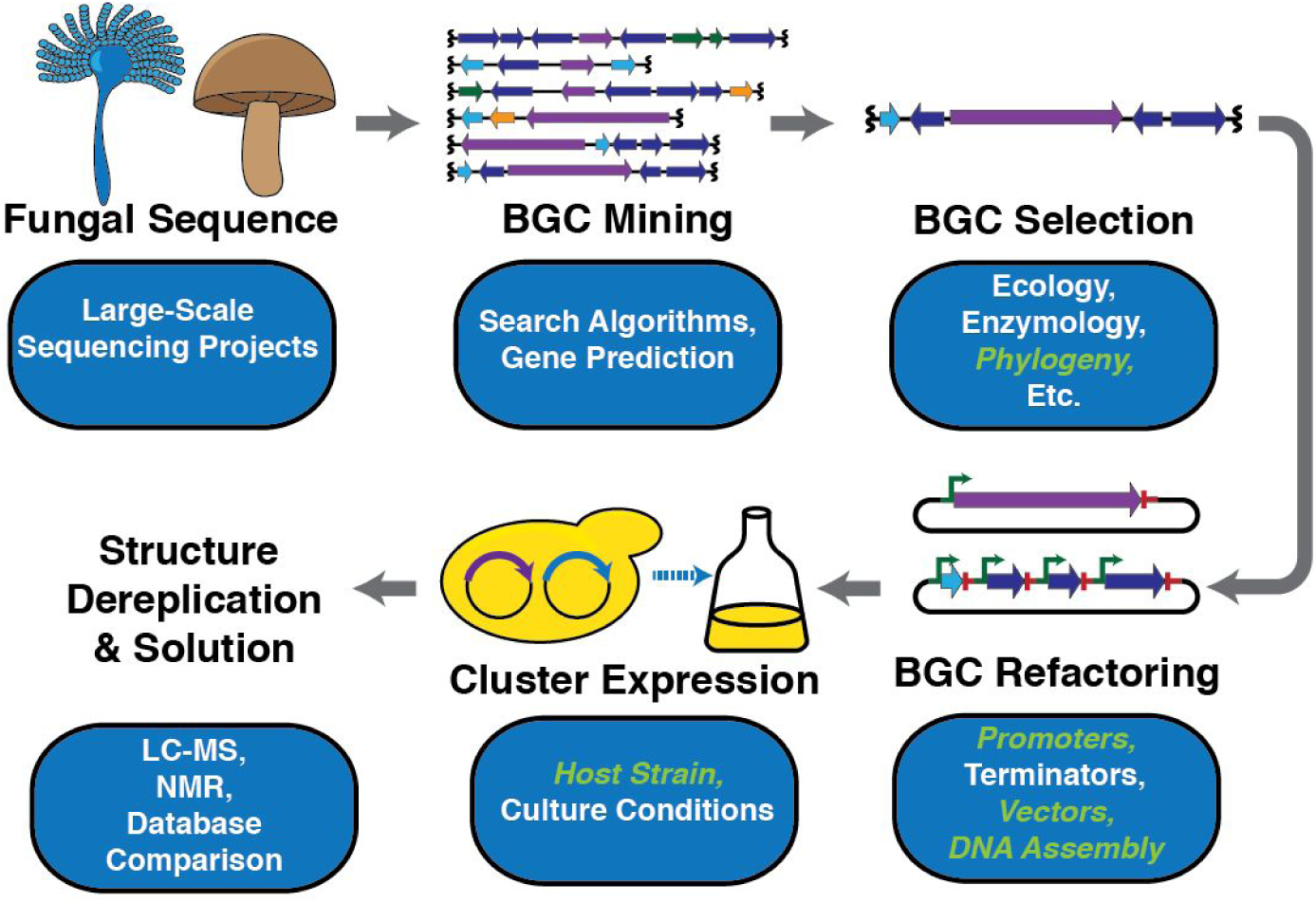
Standard workflow for heterologous expression. Aspects in green italics are addressed in this study.

**Figure 2:**
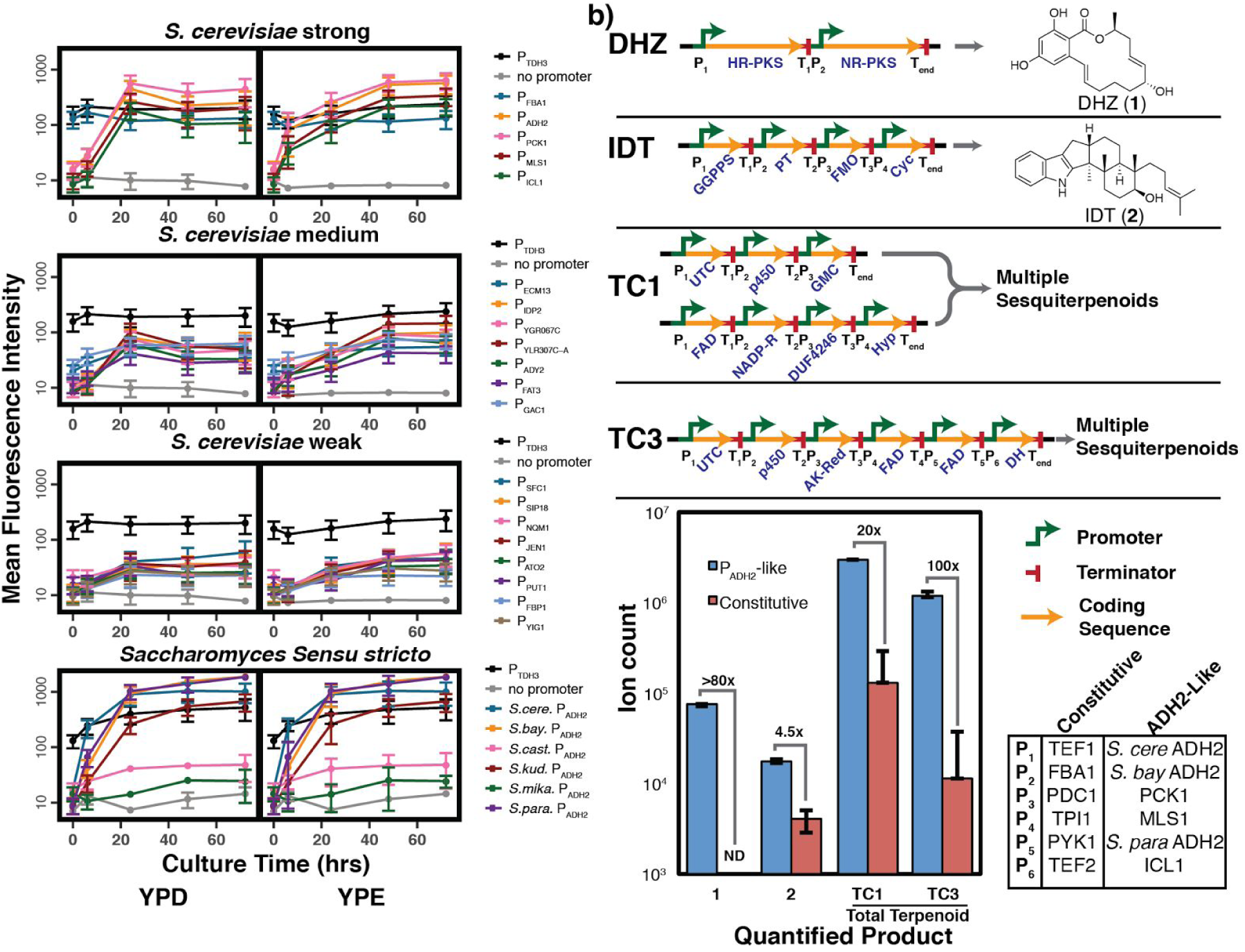
Tools developed for the HEx platform. **a)** eGFP expression from a series of P_ADH2_-like promoters in cultures grown under both fermentative (YPD) and respiratory (YPE) conditions. All fluorescence intensities are reported as the mean of three biological replicates. Error bars represent one standard deviation (n=3). **b)** Four fungal BGCs, two controls and two previously uncharacterized systems, each produce improved titers when heterologously expressed using P_ADH2_-like promoters as compared to strong constitutive promoters. ND = Not detected. Error bars represent one standard deviation (n=3). Quantitation for TC1 and TC3 was based on the sum of the integrations of extracted ion counts corresponding to the oxidized sesquiterpenoids outlined in **Supplementary Table 4.**

**Figure 3:**
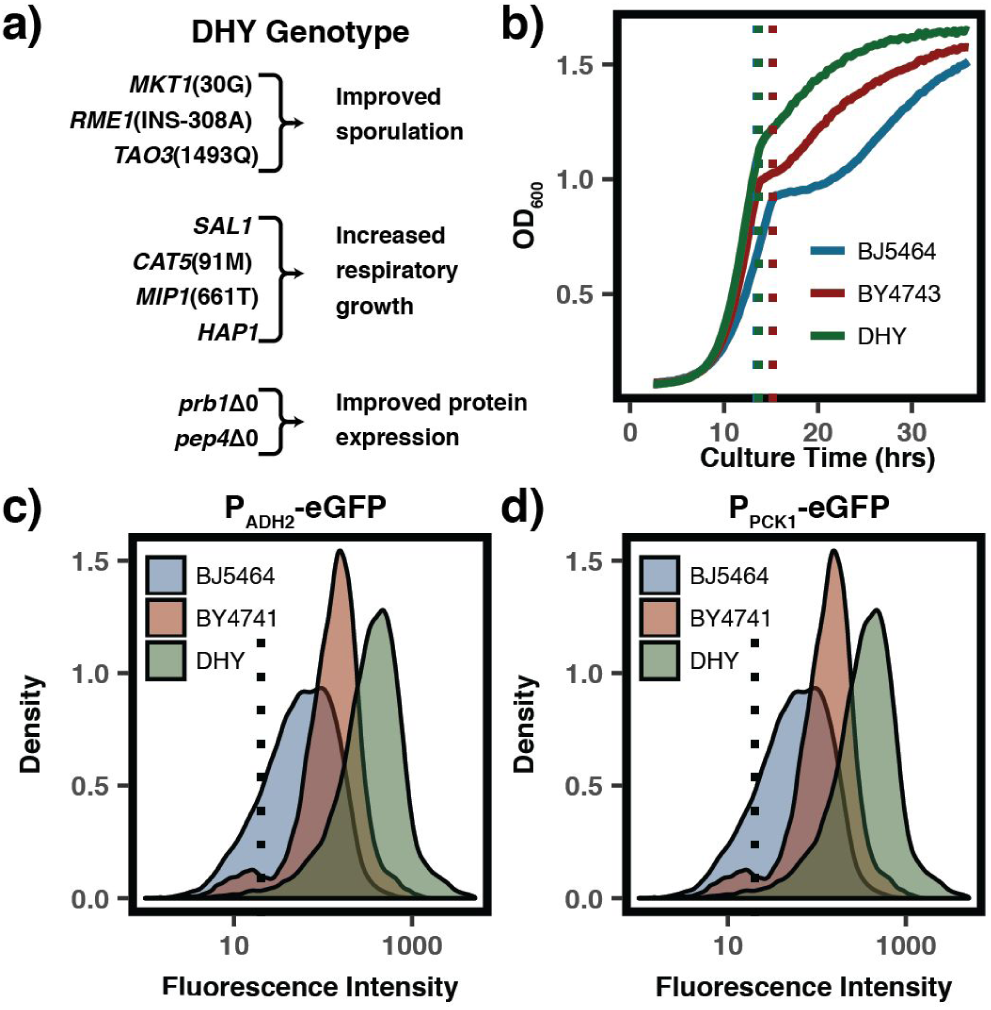
Description and characterization of DHY strains. **a)** Annotated genotype of the DHY yeast background. **b)** DHY derived yeast strain JHY702 shows improved growth, particularly after diauxic shift. Growth curves are representative of six biological replicates. Density plots for fluorescence intensity in multiple backgrounds shows significantly improved eGFP expression when driven by both **c)** P_ADH2_ and **d)** P_PCK1_. Density plots represent the fluorescence intensity of 10^4^ individual cells.

**Figure 4:**
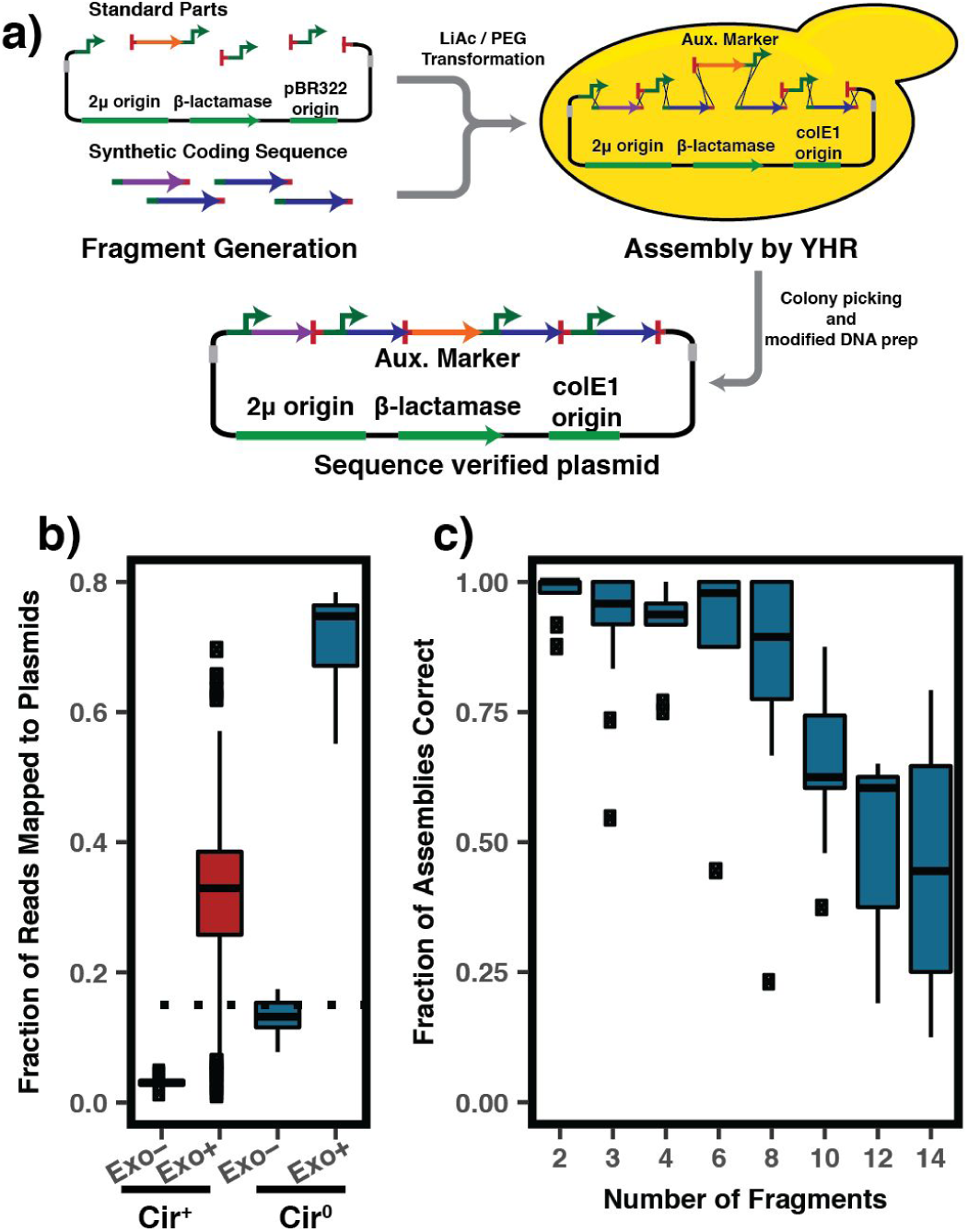
DNA assembly by yeast homologous recombination. **a)** DNA assembly from commercially synthesized fragments and genetic parts using yeast homologous recombination. **b)** Modified yeast plasmid preparation (exo+) leads to increased number of sequencing reads mapping to plasmid DNA. Dotted line marks the efficiency threshold to allow sequencing of 192 samples on a single MiSeq run. **c)** Efficient assembly of up to 14 unique DNA parts can be achieved using the protocol outlined here. Data based on 78 unique assemblies.

Previous reports of complete pathway refactoring for expression in fungal hosts have demonstrated the utility of this approach, but have been limited to the study of a single cluster(*21*–*25*). Here, we applied HEx to the expression of 41 cryptic fungal BGCs, 22 (54%) of which resulted in compounds not natively present in yeast (Figure 5-6, Table 1, **supplementary text**). The 41 BGCs were derived from diverse fungal species and include genes encoding either a membrane-bound, UbiA-like terpene cyclase(*26*) (UTCs) or a polyketide synthase (PKS) enzyme at their core. Two interesting biosynthetic insights were revealed from this study: first, UTCs represent a general class of biosynthetic enzymes present in a variety of both ascomycete and basidiomycete genomes. Second, a divergent basidiomycete clade of polyketide synthases has the unusual property of incorporating amino acids in the absence of any nonribosomal peptide synthetase (NRPS) enzymes.

**Table 1.** Summary of control and cryptic fungal BGCs examined in this study.

| ID | Type | Species of origin | Division | Productive? |
| --- | --- | --- | --- | --- |
| IDT | Ctl | *Aspergillus tubingensis* | Ascomycota | Y |
| DHZ | Ctl | *Hypomyces subiculosus* | Ascomycota | Y |
| PKS1 | PKS | *Coniothyrium sporulosum* | Ascomycota | Y |
| PKS2 | PKS | *Coniothyrium sporulosum* | Ascomycota | Y |
| PKS3 | PKS | *Acremonium* Sp. KY4917 | Ascomycota | N |
| PKS4 | PKS | *Aspergillus niger* | Ascomycota | Y |
| PKS5 | PKS | *Thielavia terrestris* | Ascomycota | N |
| PKS6 | PKS | *Trichoderma virens* | Ascomycota | Y |
| PKS7 | PKS | *Pseudogymnoascus pannorum* | Ascomycota | N |
| PKS8 | PKS | *Scedosporium apiospermum* | Ascomycota | Y |
| PKS9 | PKS | *Metarhizium anisopliae* | Ascomycota | N |
| PKS10 | PKS | *Cochliobolus heterostrophus* | Ascomycota | Y |
| PKS11 | PKS | *Pseudogymnoascus pannorum* | Ascomycota | N |
| PKS12 | PKS | *Pseudogymnoascus pannorum* | Ascomycota | N |
| PKS13 | PKS | *Pseudogymnoascus pannorum* | Ascomycota | Y |
| PKS14 | PKS | *Verruconis gallopava* | Ascomycota | Y |
| PKS15 | PKS | *Moniliophthora roreri* | Basidiomycota | Y |
| PKS16 | PKS | *Punctularia strigosozonata* | Basidiomycota | Y |
| PKS17 | PKS | *Hydnomerulius pinastri* | Basidiomycota | Y |
| PKS18 | PKS | *Arthroderma gypseum* | Ascomycota | Y |
| PKS19 | PKS | *Setosphaeria turcica* | Ascomycota | N |
| PKS20 | PKS | *Pyrenophora teres* | Ascomycota | Y |
| PKS21 | PKS | *Cladophialophora yegresii* | Ascomycota | N |
| PKS22 | PKS | *Talaromyces cellulolyticus* | Ascomycota | Y |
| PKS23 | PKS | *Endocarpon pusillum* | Ascomycota | Y |
| PKS24 | PKS | *Talaromyces cellulolyticus* | Ascomycota | Y |
| PKS25 | PKS | *Moniliophthora roreri* | Basidiomycota | N |
| PKS26 | PKS | *Hypholoma sublateritium* | Basidiomycota | N |
| PKS27 | PKS | *Ceriporiopsis subvermispora* | Basidiomycota | N |
| PKS28 | PKS | *Moniliophthora roreri* | Basidiomycota | Y |
| TC1 | UTC | *Trichoderma Virens* | Ascomycota | Y |
| TC2 | UTC | *Trichoderma Virens* | Ascomycota | N |
| TC3 | UTC | *Botryotonia cinerea* | Ascomycota | Y |
| TC4 | UTC | *Formitiporia mediterranea* | Basidiomycota | Y |
| TC5 | UTC | *Heterobasidion annosum* | Basidiomycota | Y |
| TC6 | UTC | *Gelatoporia subvermispora* | Basidiomycota | N |
| TC7 | UTC | *Dichomitus squalens* | Basidiomycota | N |
| TC8 | UTC | *Pleurotus ostreatus* | Basidiomycota | N |
| TC9 | UTC | *Schizophyllum commune* | Basidiomycota | Y |
| TC10 | UTC | *Stereum hirsutum* | Basidiomycota | N |
| TC11 | UTC | *Stereum hirsutum* | Basidiomycota | N |
| TC12 | UTC | *Dichomitus squalens* | Basidiomycota | N |
| TC13 | UTC | *Dacryopinax primogenitus* | Basidiomycota | N |
|  |  |  | **Total** | **43** |
|  |  |  | **Productive** | **24** |

Such a large-scale study of cryptic fungal natural products has not been possible until now, and was enabled through recent breakthroughs in DNA sequencing and DNA synthesis, combined with the breakthroughs in the heterologous host developed here in the HEx platform. These results reveal that the unstudied fungal sequences accumulating in genome databases can be functionally characterized in a scalable way with HEx, enabling the production of novel molecules that have never been observed in nature and paving the way for discovery of the next generation of natural products throughout the fungal kingdom.

**Figure 5:**
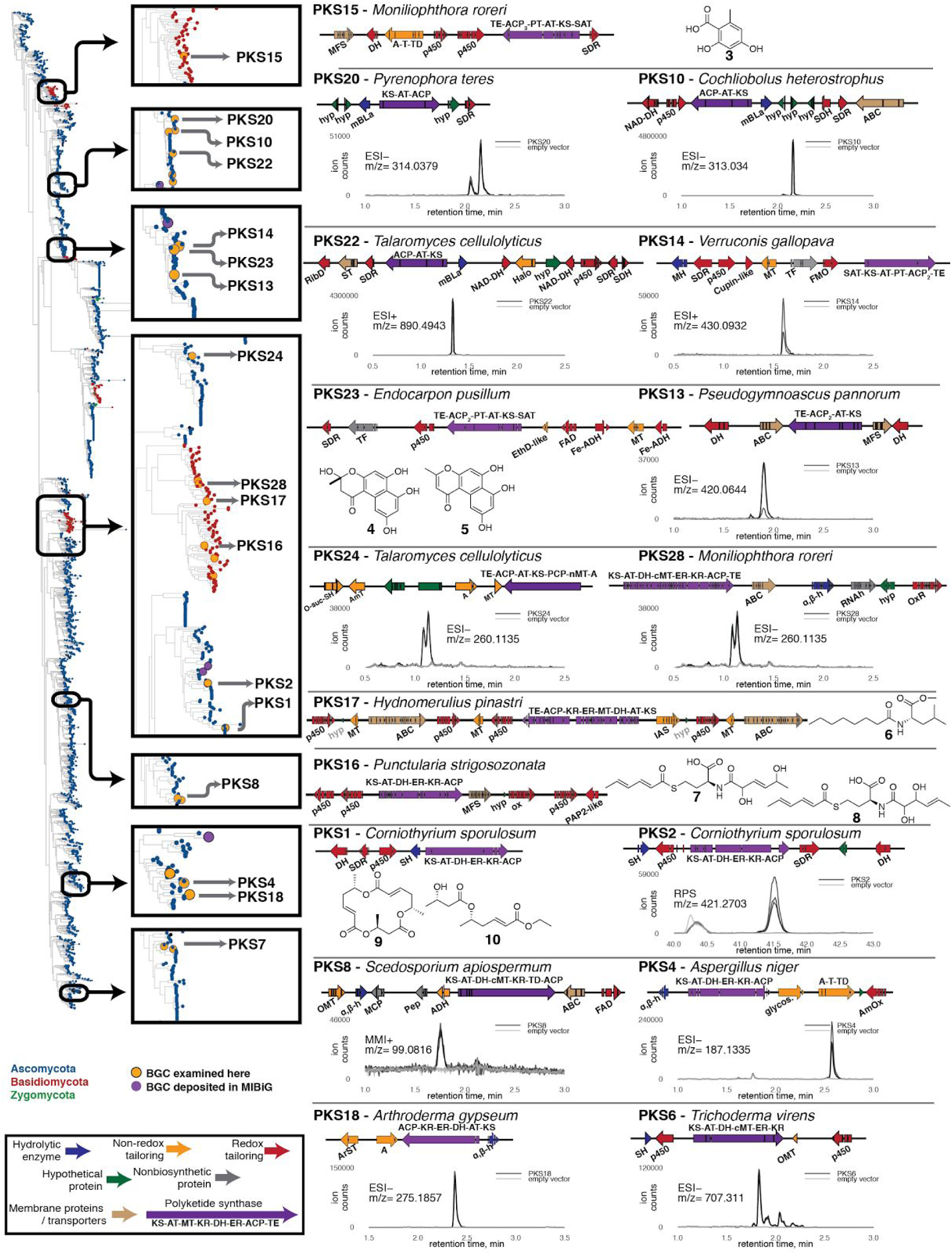
PKS BGCs examined for this study. All putative gene function abbreviations are listed in **Table S9**. Cladogram was constructed as described in the methods. All plots are the chromatograms of the specified extracted ion in three biological replicates each of both the strain expressing the BGC and an empty vector control strain. Chromatograms are data collected with electrospray ionization in either positive (ESI+), negative (ESI-), or rapid polarity switching (RPS) mode or with multi-mode ionization in positive mode (MMI). Expression strains are outlined in **Table S10** and EICs of of novel products are shown in the **Figures S7-S23**.

## Characterization of Genetic Parts

HEx is enabled by a new panel of yeast promoters that we selected and characterized to satisfy two distinct design criteria. First, the expression of BGC genes should be regulatable and coordinated. Second, each promoter sequence should be sufficiently unique so as to be compatible with homologous recombination-based DNA assembly techniques. Small panels of promoters with fewer than four members that meet these criteria have been previously reported (*27*–*30*) but were insufficient for expression of our chosen BGCs, which range from 3 to 14 genes in size.

Previous studies have demonstrated the utility of the yeast ADH2 promoter (P_ADH2_) for the heterologous expression of a variety of biosynthetic genes(*31*–*33*). P_ADH2_, which is repressed by glucose, is inactive during fermentative growth, with activation occurring only after diauxic shift. Thus, P_ADH2_ is auto-inducible(*34*) in media containing glucose and other fermentable carbon sources that are converted to non-fermentable carbon sources.

To allow for the assembly and coordinated, auto-inducible expression of entire BGCs on a small number of plasmids, we identified and characterized a panel of sequence-divergent promoters functionally similar to P_ADH2_. First, we identified 48 genes within a published *S. cerevisiae* transcriptome dataset(*35*) that appeared co-regulated with ADH2 in that their transcripts were weakly expressed in mid-log phase fermentative growth, but were highly abundant during respiration (**Table S2**). We then constructed single-copy chromosomal integrations of the corresponding promoters driving expression of a GFP gene and measured fluorescence. Four promoters (P_ADH2_, P_MLS1_, P_PCK1_, P_ICL2_) demonstrated the desired delayed-induction phenotype with expression levels after 24 hours being similar to or greater than P_TDH3_ or P_FBA1_, two commonly-used strong constitutive promoters (Figure 2a)(*36*, *37*). We also identified 23 promoters that were co-regulated with P_ADH2_, but whose degree of induction was slightly or significantly lower, thus providing opportunities for lower levels of coordinated gene expression (Figure 2a, **Figure S2a-c, Table S3**). Among these 20 promoters, no two have greater than 29% identity, demonstrating that these sequences are sufficiently unique to allow assembly using homology-based approaches. Additionally, we characterized the promoters of the closest *ADH2* homologue in each of five close relatives of *S. cerevisiae*, the *sensu stricto Saccharomyces* species (**Figure S1d-f**). Three of these promoters (from *S. paradoxus, S. bayanus*, and *S. kudriavzevii*) were functionally equivalent to the S. cerevisiae homologue (Figure 2a), bringing the number of P_ADH2_-like promoters to 30, with 7 having strengths at least equal to strong constitutive promoters. We refer to these P_ADH2_-like promoters as the HEx promoters.

To study the utility of these promoters for BGC engineering, we chose to engineer versions of four BGCs on 2 µm plasmids with expression of each gene driven by a either a strong HEx promoter or a strong constitutive promoter. Two of these clusters, DHZ and IDT, were controls, selected as they are known to function in yeast and are the producers of the polyketide 7’,8’-dehydrozearalenol (DHZ, **1**) and the indole diterpene **2** (IDT), respectively. Additionally, we selected two uncharacterized BGCs containing a UbiA-type sesquiterpene cyclase (UTC, Figure 2b, TC1, TC3). Analysis of the control clusters demonstrated that production of compound **1** was detectable only when expression was driven by the HEx promoters, and undetectable with constitutive promoters. Titers of **2** were 4.5-fold higher with HEx promoters than with constitutive expression. For the uncharacterized UTC containing clusters, combined titers for the oxygenated sesquiterpenoids (**Table S5**) produced by both clusters were significantly greater (20-fold for TC1, 100-fold for TC3) when refactored with P_ADH2_-like sequences versus strong constitutive promoters. These results establish the broad utility of P_ADH2_-like, delayed-induction HEx promoters as tools for the coordinated expression of multiple heterologous proteins in yeast.

## Improved Host Strains

HEx promoters are active only under respiratory conditions, which necessitates their use in yeast host strains with functional mitochondria. We chose to use the well-characterized S288c-derived strains BY4741 and BY4742(*38*) as our starting point for host optimization. These and related strains have been used in previous heterologous expression studies in yeast with great success(*17*, *39*) despite known mitochondrial genome stability defects present in all strains in the S288c lineage, as indicated by increased petite frequencies(*40*). The alleles leading these traits have been reported(*40*, *41*), so to better facilitate the heterologous expression of fungal BGCs in yeast, we generated strains in which these deficiencies, as well as vestigial defects in sporulation efficiency present in S288c-derived strains, were repaired.

These defects were repaired in an improved strain background named DHY (Figure 3a). Alleles absent from all S288c derived strains that lead to increased mitochondrial stability (*SAL1 CAT5*(91M) *MIP1*(661T) *HAP1*)(*40*, *41*) and high sporulation (*MKT1*(30G) *RME1*(INS-308A) *TAO3*(1493Q)) were introduced(*42*). These alterations increased sporulation from 2% to 62% after two days while decreasing petite frequency from 52% to 2.5%. Additionally, we deleted the *PEP4* and *PRB1* vacuolar protease encoding genes as in BJ5464, a strain with demonstrated improvements in heterologous protein production(*43*). We generated a panel of strains, both prototrophic and auxotrophic, of both mating types (*MAT* **a** and *MAT* **α**), with these nine beneficial changes. For BGC expression, we also integrated several genes for essential post-translational modification enzymes. npgA, a *holo*-ACP synthase from *Aspergillus nidulans* that has demonstrated flexibility for generation of a variety of holo-carrier proteins for both PKS(*39*) and nonribosomal peptide synthetase (NRPS)(*44*) containing systems expressed in *S. cerevisiae*. Additionally, as cytochrome P450s are ubiquitous in fungal BGCs, ATEG_05064, a cytochrome P450 reductase *Aspergillus terreus* has been engineered into our strain background. The full panel of strains used in this study is listed in **Table S4**.

The engineered DHY background exhibits improved respiratory growth as compared to both BY4741 and BJ5464 (Figure 3b). GFP expression driven by P_ADH2_ (Figure 3c) and P_PCK1_ (Figure 3d) also showed marked improvement in DHY. Not only was the mean expression significantly increased, but a population of non-fluorescent cells prevalent in the BY4741 culture were undetectable in the DHY-derived strain, likely a result of the improved mitochondrial function during expression-inducing respiratory growth conditions.

Compared to commonly used lab strains, the improved genetic tractability, growth, and expression characteristics of the DHY background make it an ideal host strain for the HEx platform and for heterologous protein expression more generally.

## High-throughput DNA Assembly

Heterologous expression of microbial BGCs necessitates a high-throughput, low-cost means of assembling large, multigene constructs expressing cryptic BGCs. Yeast homologous recombination represents such an approach and has been previously applied to the refactoring of large BGCs for expression in model bacterial hosts utilizing intact DNA from native producing strains or environmental samples(*45*–*49*). In the absence of this native DNA to be used as a PCR template, BGC sequence must be sourced from commercial vendors. At present, commercial providers of synthetic DNA supply gene-sized fragments at a relatively low cost, but do not offer cost-effective solutions for the larger DNAs required for studying BGCs, which are often >20 kb. Additionally, synthesis of AT-rich yeast regulatory sequences, especially promoters, has proven challenging for commercial providers. We therefore purchased synthetic DNA encoding the protein-coding regions of BGC genes, and developed an improved means to assemble these genes into large multigene cassettes including promoters, terminators, and expression vectors.

The strategy employed here for cluster refactoring used to design parts for assembly by yeast homologous recombination is illustrated in Figure 4a. DNAs for adjacent fragments were designed with 50 bp of overlapping sequence. In cases where a gene was small enough to be ordered as a single DNA fragment, overlapping sequence to both the flanking promoter and terminator were added, whereas with genes that were split into multiple DNA fragments, overlapping sequences were added to adjacent sequences. All clusters were refactored by building plasmids of 7 or fewer genes each. Each gene within a plasmid was flanked by promoters and terminators used in the order defined in **Table S6**. Placing overlapping sequences exclusively on the coding sequence fragments allowed for the same standard parts (promoters, terminators, and linearized vectors) to be generated in bulk and used in all assemblies (**Table S7**). For assemblies involving three or more genes, an auxotrophic marker was placed between the second terminator and third promoter with no marker present on the vector. By applying the constraint of the auxotrophic marker and origin of replication being on separate fragments, assembly of incorrect plasmids was significantly reduced.

Most previous YHR-based assembly techniques relied on passage of vectors through *E. coli* to generate large amounts of DNA(*50*). We developed a protocol for the purification of plasmid DNA directly from yeast clones wherein the majority of contaminating yeast genomic DNA was removed by exonuclease treatment. This procedure enabled high-throughput DNA sequencing libraries to be prepared directly from yeast colonies, simplifying the process of verifying that target plasmids were correctly assembled. Unmapped sequencing reads in exonuclease treated samples mapped primarily to the native 2µm plasmid present in the majority of laboratory yeast strains (cir^+^ strains). this hypothesis was confirmed by demonstrating that sequencing plasmid DNA out of Y800(*51*), a strain lacking this plasmid (cir^0^), led to greater than 75% of reads mapping to the desired plasmid (Figure 4b).

The HEx process is a simplified workflow with increased throughput and decreased cost relative to other YHR techniques. We found that assemblies of up to 14 unique DNA fragments can routinely be achieved with high efficiency (Figure 4c). Overall, we have applied HEx to assemble 41 gene clusters and sequence >1000 yeast clones.

## Expression of Cryptic BGCs

To apply the HEx platform on a large scale, we chose to examine two classes of fungal BGCs: those encoding either a polyketide synthase (PKS) or a UbiA-type sesquiterpene cyclase (UTC) as their core enzyme. Phylogenetic analysis has suggested that much chemical diversity remains to be discovered in fungal PKSs, with the possible existence of entirely unstudied classes(*52*). UTCs represent a newly discovered class of membrane-bound terpene cyclases, homologous to UbiA prenyltransferases, discovered during the recent elucidation of the biosynthesis of fumagillin(*53*).

We developed a computational pipeline to prioritize PKS- and UTC-containing BGCs for expression with HEx. We studied all 581 sequenced fungal genomes publicly available in the GenBank database of the National Center for Biotechnology Information (NCBI, as of July 2015). We analyzed each genome for BGCs using antiSMASH2(*54*), identifying 3512 BGCs harboring an iterative type 1 PKS and 326 BGCs harboring a UTC homologue. We generated phylogenetic trees of each of these enzyme types, identified characterized homologs from the MIBiG database(*55*), and selected BGCs from clades having few characterized members (Figure 5, Figure 6). These BGCs were found in the genomes of both ascomycetes and basidiomycetes. Basidiomycetes have historically been more difficult than ascomycetes to culture with fewer tools for genetic manipulation available(*56*). As a result, BGCs from basidiomycetes are under-studied, with only two PKS-containing clusters deposited in MIBig as of writing, suggesting that these organisms represent a reservoir of under-studied BGCs. All coding sequences were ordered as a series of fused exons with no codon optimization unless required for DNA synthesis (on average approximately 1 change per 5000 bp). The start codons, stop codons, and intron/exon boundaries were exactly as deposited in GenBank (**Table S8**).

**Figure 6:**
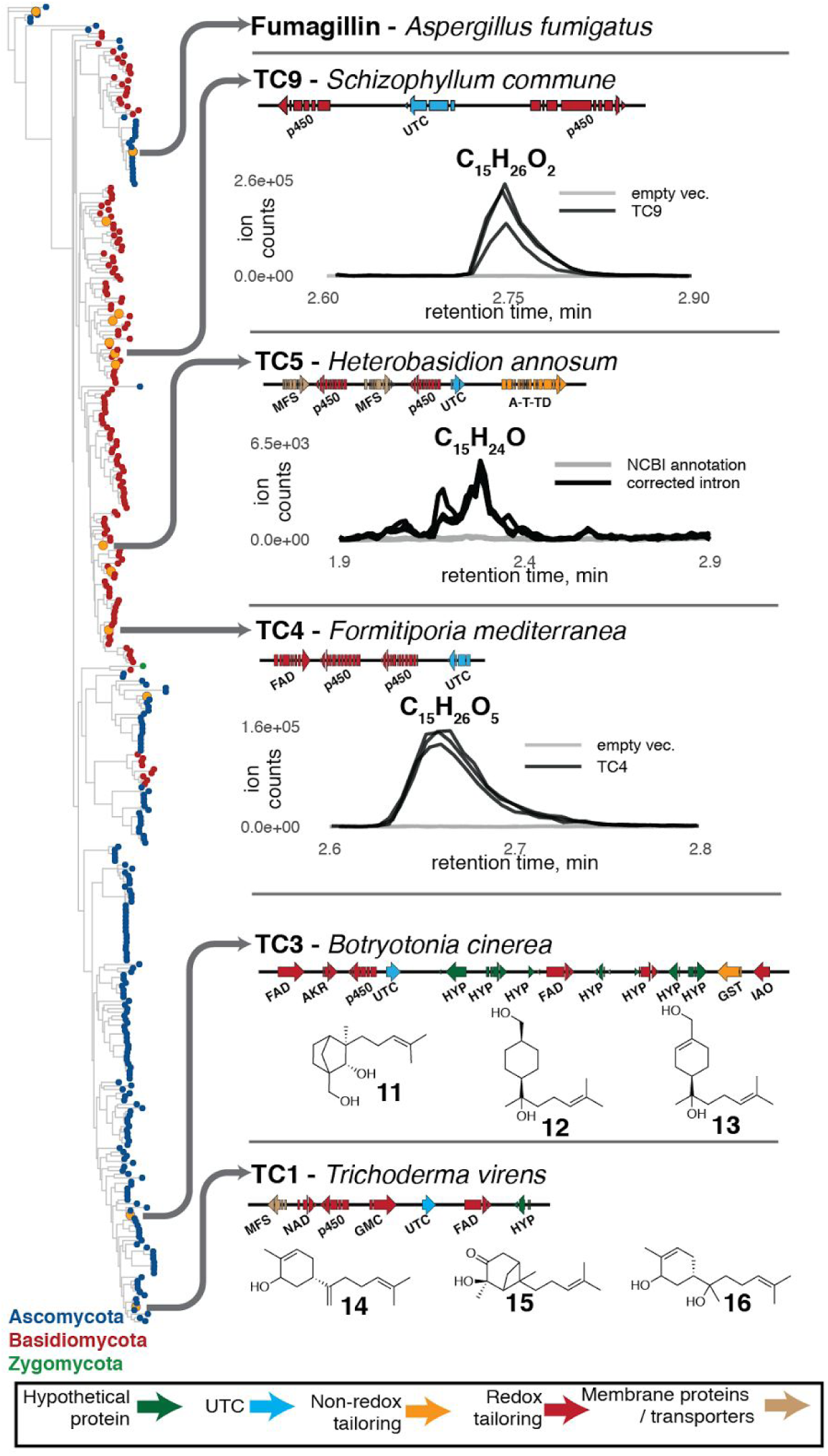
UbiA-type cyclases represent a general class of biosynthetic enzymes. Putative enzyme activity abbreviations are listed in **Table S9**. Cladogram generated using UTC cyclase sequence. The cyclases associated with all clusters examined in this study are denoted by orange tips in the cladogram.

In order to explore novel fungal PKSs, we began with the hypothesis that novel PKS sequence would lead to novel compounds. In order to select unusual PKS BGCs, we performed phylogenetic analysis of the ketosynthase sequences of all 3512 PKS sequences found in the 581 sequenced fungal genomes (Figure 5, **Figure S7**). We first identified sequences that existed in clades where few or no characterized BGCs were found. To further narrow the list to BGCs likely to produce a compound, we selected those whose genetic structure was conserved across 3 or more species, and that contained an in-*cis* or proximal in-*trans* protein capable of releasing the polyketide from the carrier protein of the PKS (**Figure S4**). From the BGCs that met these criteria, we selected 28, containing between 3 and 11 genes, for characterization with HEx. Seven of these were derived from basidiomycete-specific clades (Figure 5) while the remaining 21 were found in the genomes of Ascomycetes.

Of the 7 basidiomycete BGCs chosen, three (PKS16, PKS17, and PKS28) produced natural products. The production of 7 and 8 from PKS16, both of which are novel N-, S- bis-acylated amino acids, is unprecedented, as they incorporate an amino acid but the cluster contains no NRPS gene (Figure 5, **Figure S4**). Similarly, PKS17 produces compound 6, a leucine O-methyl ester with an additional polyketide chain amidated to the amino ester. PKS28 produced a pair of compounds that were not structurally characterized, but on the basis of high-resolution mass spectral data, are likely to contain at least one nitrogen atom. To our knowledge, these are the first examples of fungal BGCs producing polyketide-amino acid hybrid compounds in the absence of NRPS encoding genes.

Of the 21 ascomycete-derived PKS clusters, 13 produced compounds. The most notable was the PKS1 cluster, which only contained an PKS, a hydrolase, and the genes for three tailoring enzymes: a cytochrome P450 monooxygenase, a flavin-dependent monooxygenase (FMO), and a short-chain reductase (SDR) (**Table S9**). This cluster produced **9** and **10** as major products (Figure 5) along with a variety of oligo-esters. **9**, an asymmetric macrotriolide, results from the condensation of two triketides with a single diketide and closely resembles the macrosphelide family of fungal natural products, compounds with antimicrobial activity(*57*) whose BGCs have yet to be elucidated.

In addition to these novel compounds, two clusters produced known compounds and novel derivatives thereof. PKS15 produced orsellinic acid (**3**) as the major product along with several other higher molecular weight compounds. PKS23 produced **4** and **5** as major products, along with several additional putative products of higher mass. **4** and **5** are both precursors of phenalenone, a compound whose biosynthetic gene cluster was elucidated after the selection of PKS23 for expression in this study(*58*). Taken together, these results demonstrate the power of the HEx platform to produce both novel and previously known compounds using unstudied BGCs derived from uncultivated fungi.

To study fungal UTCs, we constructed a phylogenetic tree based on the UbiA-type sesquiterpene cyclase, Fma-TC, from the fumagillin biosynthetic pathway(52) (Figure 6). The cytochrome P450 monoxygenase, Fma-P450, from the fumagillin pathway is a powerful enzyme catalyzing the 8*e* oxidation of bergamotene to generate a highly oxygenated product(*9*). We selected 13 UTC-containing BGCs spanning the entirety of the cladogram in Figure 6 where a cytochrome P450 monooxygenase gene was proximal to the UTC gene (**Figure S5a**). Screening of strains expressing these clusters by LC/HRMS revealed novel spectral features consistent with oxidized sesquiterpenoids produced by five clusters (Figure 4). The structures of the major compounds produced by TC1 (compounds **14**, **15**, and **16**) and TC3 (compounds **11**, **12**, and **13**) were elucidated by NMR (**supplementary text**). Unique among these clusters is TC9 from the basidiomycete *Schizophyllum commune* where the UTC alone produces a series of sesquiterpenols that, when placed in the context of the full cluster, are further oxidized by the two adjacent cytochrome P450 monooxygenases. These results not only demonstrate a series of structurally novel sesquiterpenoids, they also demonstrate that the membrane-bound UTCs represent a general class of terpene cyclase encoded by the genomes of diverse fungi.

Including both PKS and UTC BGCs, we found that 19 of the 41 clusters studied did not produce detectable compounds. We hypothesized that gene annotation errors introduced by incorrect intron prediction was likely to be a common failure mode in the expression of cryptic fungal BGCs, and therefore sought to rescue production by improved intron annotation. Manual inspection of one UTC (TC5) that had yielded no products suggested an incorrect intron prediction at the 5’ terminus of the gene (**Figure S5b**). Correction of this intron led to a C-terminal protein sequence that aligned well with known functional UTCs. When tested in the HEx pipeline, the version with the corrected intron produced an oxidized sesquiterpenoid (Figure 6) confirming that incorrect intron prediction can be a failure mode in approaches that rely on publicly available gene annotations. These results illustrate the importance of careful gene curation and the need for improved eukaryotic gene prediction, particularly with sequences from taxa with few studied members. We anticipate this being particularly important for BGCs derived from basidiomycetes as introns are more common in this phyla than in filamentous ascomycetes(*59*).

In conclusion, using the HEx platform developed here, we built strains expressing 41 cryptic fungal BGCs. 22 (54%) of these clusters, derived from diverse ascomycete and basidiomycete fungal species, produced detectable levels of compounds not native to *S. cerevisiae* (Table 1). Ongoing and future studies will work to improve this success rate through a detailed analysis of those BGCs that failed. Testing multiple splice variants for ambiguous genes, quantifying transcript and protein expression levels for each gene, and ensuring phosphopantetheinylation of all ACP domains are among the approaches that may provide insight into the common failure modes of fungal BGCs refactored for expression in yeast. Additionally, varying protein stoichiometry through building multiple versions of each refactored cluster with varying promoter strengths may also resurrect nonfunctional clusters or increase conversion of biosynthetic intermediates in those that produce multiple products.

A recent analysis of the diversity of natural products discovered over time has highlighted the need for innovative new approaches for molecule discovery(*60*). Here, by performing a large-scale survey of diverse BGCs from across the fungal kingdom, we have demonstrated such an approach. Using our platform, we identified a panel of novel natural products produced by enzymes with novel activities. Moreover, the genetic parts, improved host strains, and DNA assembly pipeline that comprise the HEx platform provide an improved means for accessing the vast biosynthetic potential encoding natural products with novel structures and bioactivities that exist within the multitude of cryptic BGCs present in fungal genomes.

## Acknowledgements

CJBH, BN, YT, MM and MEH all have a financial interest in Hexagon Bio Inc.

## Supplementary Materials

Materials and Methods

Supplementary Text

Figures S1-S78

Tables S1-S12

Supplementary Sequences

## References and Notes

1. D. J. Newman, G. M. Cragg, Natural Products as Sources of New Drugs from 1981 to 2014. J. Nat. Prod. 79, 629–661 (2016).

2. A. Schueffler, T. Anke, Fungal natural products in research and development. Nat. Prod. Rep. 31, 1425–1448 (2014).

3. P. Wiemann, N. P. Keller, Strategies for mining fungal natural products. J. Ind. Microbiol. Biotechnol. 41, 301–313 (2014).

4. M. Blackwell, The Fungi: 1, 2, 3 & 5.1 million species? Am. J. Bot. 98, 426–438 (2011).

5. D. O. Inglis et al., Comprehensive annotation of secondary metabolite biosynthetic genes and gene clusters of Aspergillus nidulans, A. fumigatus, A. niger and A. oryzae. BMC Microbiol. 13, 91 (2013).

6. N. Khaldi et al., SMURF: Genomic mapping of fungal secondary metabolite clusters. Fungal Genet. Biol. 47, 736–741 (2010).

7. M. Keller, K. Zengler, Tapping into microbial diversity. Nat. Rev. Microbiol. 2, 141–150 (2004).

8. A. A. Brakhage, Regulation of fungal secondary metabolism. Nat. Rev. Microbiol. 11, 21–32 (2012).

9. H.-C. Lin et al., Generation of complexity in fungal terpene biosynthesis: discovery of a multifunctional cytochrome P450 in the fumagillin pathway. J. Am. Chem. Soc. 136, 4426–4436 (2014).

10. S. Bergmann et al., Activation of a silent fungal polyketide biosynthesis pathway through regulatory cross talk with a cryptic nonribosomal peptide synthetase gene cluster. Appl. Environ. Microbiol. 76, 8143–8149 (2010).

11. H.-H. Yeh et al., Resistance Gene-Guided Genome Mining: Serial Promoter Exchanges in Aspergillus nidulans Reveal the Biosynthetic Pathway for Fellutamide B, a Proteasome Inhibitor. ACS Chem. Biol. 11, 2275–2284 (2016).

12. J. Weber et al., Functional Reconstitution of a Fungal Natural Product Gene Cluster by Advanced Genome Editing. ACS Synth. Biol. 6, 62–68 (2017).

13. D. A. Adpressa, K. J. Stalheim, P. J. Proteau, S. Loesgen, Unexpected Biotransformation of the HDAC Inhibitor Vorinostat Yields Aniline-Containing Fungal Metabolites. ACS Chem. Biol. (2017), doi: 10.1021/acschembio.7b00268.

14. M. T. Henke et al., New Aspercryptins, Lipopeptide Natural Products, Revealed by HDAC Inhibition in Aspergillus nidulans. ACS Chem. Biol. 11, 2117–2123 (2016).

15. X.-M. Mao et al., Epigenetic genome mining of an endophytic fungus leads to the pleiotropic biosynthesis of natural products. Angew. Chem. Int. Ed Engl. 54, 7592–7596 (2015).

16. K. D. Clevenger et al., A scalable platform to identify fungal secondary metabolites and their gene clusters. Nat. Chem. Biol. (2017), doi: 10.1038/nchembio.2408.

17. D.-K. Ro et al., Production of the antimalarial drug precursor artemisinic acid in engineered yeast. Nature. 440, 940–943 (2006).

18. S. Galanie, K. Thodey, I. J. Trenchard, M. Filsinger Interrante, C. D. Smolke, Complete biosynthesis of opioids in yeast. Science. 349, 1095–1100 (2015).

19. S. Brown, M. Clastre, V. Courdavault, S. E. O’Connor, De novo production of the plant-derived alkaloid strictosidine in yeast. Proc. Natl. Acad. Sci. U. S. A. 112, 3205–3210 (2015).

20. C. A. Smith, E. J. Want, G. O’Maille, R. Abagyan, G. Siuzdak, XCMS: processing mass spectrometry data for metabolite profiling using nonlinear peak alignment, matching, and identification. Anal. Chem. 78, 779–787 (2006).

21. Y.-M. Chiang et al., An Efficient System for Heterologous Expression of Secondary Metabolite Genes in Aspergillus nidulans. J. Am. Chem. Soc. 135, 7720–7731 (2013).

22. M. N. Heneghan et al., First heterologous reconstruction of a complete functional fungal biosynthetic multigene cluster. Chembiochem. 11, 1508–1512 (2010).

23. T. Itoh, T. Kushiro, I. Fujii, in Fungal Secondary Metabolism (2012), pp. 175–182.

24. M. T. Nielsen et al., Heterologous reconstitution of the intact geodin gene cluster in Aspergillus nidulans through a simple and versatile PCR based approach. PLoS One. 8, e72871 (2013).

25. D. J. Smith, M. K. R. Burnham, J. Edwards, A. J. Earl, G. Turner, Cloning and Heterologous Expression of the Penicillin Biosynthetic Gene Cluster from Penicillium chrysogenum. Nat. Biotechnol. 8, 39–41 (1990).

26. H.-C. Lin et al., The fumagillin biosynthetic gene cluster in Aspergillus fumigatus encodes a cryptic terpene cyclase involved in the formation of β-trans-bergamotene. J. Am. Chem. Soc. 135, 4616–4619 (2013).

27. S. Labbé, D. J. Thiele, Copper ion inducible and repressible promoter systems in yeast. Methods Enzymol. 306, 145–153 (1999).

28. V. Rönicke, W. Graulich, D. Mumberg, R. Müller, M. Funk, in Methods in Enzymology (1997), pp. 313–322.

29. M. Johnston, R. W. Davis, Sequences that regulate the divergent GAL1-GAL10 promoter in Saccharomyces cerevisiae. Mol. Cell. Biol. 4, 1440–1448 (1984).

30. K. Weinhandl, M. Winkler, A. Glieder, A. Camattari, Carbon source dependent promoters in yeasts. Microb. Cell Fact. 13, 5 (2014).

31. K. M. Lee, N. A. DaSilva, Evaluation of the Saccharomyces cerevisiae ADH2 promoter for protein synthesis. Yeast. 22, 431–440 (2005).

32. C. D. Reeves, Z. Hu, R. Reid, J. T. Kealey, Genes for the biosynthesis of the fungal polyketides hypothemycin from Hypomyces subiculosus and radicicol from Pochonia chlamydosporia. Appl. Environ. Microbiol. 74, 5121–5129 (2008).

33. Y.-H. Chooi, Y. J. Hong, R. A. Cacho, D. J. Tantillo, Y. Tang, A cytochrome P450 serves as an unexpected terpene cyclase during fungal meroterpenoid biosynthesis. J. Am. Chem. Soc. 135, 16805–16808 (2013).

34. F. W. Studier, F. William Studier, Protein production by auto-induction in high-density shaking cultures. Protein Expr. Purif. 41, 207–234 (2005).

35. Z. Xu et al., Bidirectional promoters generate pervasive transcription in yeast. Nature. 457, 1033–1037 (2009).

36. J. Sun et al., Cloning and characterization of a panel of constitutive promoters for applications in pathway engineering in Saccharomyces cerevisiae. Biotechnol. Bioeng. 109, 2082–2092 (2012).

37. M. E. Lee, W. C. DeLoache, B. Cervantes, J. E. Dueber, A Highly Characterized Yeast Toolkit for Modular, Multipart Assembly. ACS Synth. Biol. 4, 975–986 (2015).

38. C. B. Brachmann et al., Designer deletion strains derived from Saccharomyces cerevisiae S288C: a useful set of strains and plasmids for PCR-mediated gene disruption and other applications. Yeast. 14, 115–132 (1998).

39. C. Bond, Y. Tang, L. Li, Saccharomyces cerevisiae as a tool for mining, studying and engineering fungal polyketide synthases. Fungal Genet. Biol. 89, 52–61 (2016).

40. L. N. Dimitrov, R. B. Brem, L. Kruglyak, D. E. Gottschling, Polymorphisms in multiple genes contribute to the spontaneous mitochondrial genome instability of Saccharomyces cerevisiae S288C strains. Genetics. 183, 365–383 (2009).

41. A. M. Deutschbauer, R. W. Davis, Quantitative trait loci mapped to single-nucleotide resolution in yeast. Nat. Genet. 37, 1333–1340 (2005).

42. J. D. Smith et al., A method for high-throughput production of sequence-verified DNA libraries and strain collections. Mol. Syst. Biol. 13, 913 (2017).

43. E. W. Jones, Tackling the protease problem in Saccharomyces cerevisiae. Methods Enzymol. 194, 428–453 (1991).

44. R. A. Cacho, Y. Tang, Reconstitution of Fungal Nonribosomal Peptide Synthetases in Yeast and In Vitro. Methods Mol. Biol. 1401, 103–119 (2016).

45. Y. Luo et al., Activation and characterization of a cryptic polycyclic tetramate macrolactam biosynthetic gene cluster. Nat. Commun. 4, 2894 (2013).

46. K. Yamanaka et al., Direct cloning and refactoring of a silent lipopeptide biosynthetic gene cluster yields the antibiotic taromycin A. Proc. Natl. Acad. Sci. U. S. A. 111, 1957–1962 (2014).

47. H.-S. Kang, Z. Charlop-Powers, S. F. Brady, Multiplexed CRISPR/Cas9- and TAR-Mediated Promoter Engineering of Natural Product Biosynthetic Gene Clusters in Yeast. ACS Synth. Biol. 5, 1002–1010 (2016).

48. O. Bilyk, O. N. Sekurova, S. B. Zotchev, A. Luzhetskyy, Cloning and Heterologous Expression of the Grecocycline Biosynthetic Gene Cluster. PLoS One. 11, e0158682 (2016).

49. Z. Feng, J. H. Kim, S. F. Brady, Fluostatins produced by the heterologous expression of a TAR reassembled environmental DNA derived type II PKS gene cluster. J. Am. Chem. Soc. 132, 11902–11903 (2010).

50. Z. Shao, H. Zhao, H. Zhao, DNA assembler, an in vivo genetic method for rapid construction of biochemical pathways. Nucleic Acids Res. 37, e16–e16 (2008).

51. N. Burns et al., Large-scale analysis of gene expression, protein localization, and gene disruption in Saccharomyces cerevisiae. Genes Dev. 8, 1087–1105 (1994).

52. Y. F. Li et al., Comprehensive curation and analysis of fungal biosynthetic gene clusters of published natural products. Fungal Genet. Biol. 89, 18–28 (2016).

53. H.-C. Lin et al., The fumagillin biosynthetic gene cluster in Aspergillus fumigatus encodes a cryptic terpene cyclase involved in the formation of β-trans-bergamotene. J. Am. Chem. Soc. 135, 4616–4619 (2013).

54. K. Blin et al., antiSMASH 2.0--a versatile platform for genome mining of secondary metabolite producers. Nucleic Acids Res. 41, W204–12 (2013).

55. M. H. Medema et al., Minimum Information about a Biosynthetic Gene cluster. Nat. Chem. Biol. 11, 625–631 (2015).

56. M. Stadler, D. Hoffmeister, Fungal natural products-the mushroom perspective. Front. Microbiol. 6, 127 (2015).

57. M. Hayashi et al., Macrosphelide, a novel inhibitor of cell-cell adhesion molecule. I. Taxonomy, fermentation, isolation and biological activities. J. Antibiot.. 48, 1435–1439 (1995).

58. S.-S. Gao et al., Phenalenone Polyketide Cyclization Catalyzed by Fungal Polyketide Synthase and Flavin-Dependent Monooxygenase. J. Am. Chem. Soc. 138, 4249–4259 (2016).

59. D. M. Kupfer et al., Introns and splicing elements of five diverse fungi. Eukaryot. Cell. 3, 1088–1100 (2004).

60. C. R. Pye, M. J. Bertin, R. S. Lokey, W. H. Gerwick, R. G. Linington, Retrospective analysis of natural products provides insights for future discovery trends. Proc. Natl. Acad. Sci. U. S. A. 114, 5601–5606 (2017).

